# Reproductive experience promotes permanent body growth independently of growth hormone

**DOI:** 10.64898/2026.04.30.721916

**Authors:** Gabriel O. de Souza, Willian O. dos Santos, Frederick Wasinski, Ligia M. M. de Sousa, Andressa G. Amaral, Daniela O. Gusmao, Edward O. List, John J. Kopchick, Gimena Fernandez, Mario Perelló, Carla R. P. Oliveira, Manuel H. Aguiar-Oliveira, Jose Donato

## Abstract

Pregnancy leads to many adaptations in the maternal body, some of which are reversible. However, reproductive experience can also result in permanent effects. Here, we investigated how pregnancy influences the somatotrophic system and the lasting effects of reproductive experience on the maternal organism. Reproductive experience induced a pronounced increase in lean body mass and longitudinal growth in both wild-type and growth hormone (GH)-deficient mice compared with age-matched virgins. Body growth was primarily observed during the first pregnancy, whereas a second gestation was mostly associated with increased adiposity. Increased GH secretion was observed in pregnant wild-type mice but not in pregnant GHRHR-deficient mice. Pregnancy-induced body growth is preserved despite disruption of GH-, ghrelin-, and estrogen-related signaling pathways. Data from a cohort of women with isolated GH deficiency (IGHD) caused by a loss-of-function mutation in the GHRHR gene revealed that nulliparous women were 7 cm shorter than those with one or more pregnancies. In conclusion, reproductive experience induces permanent changes in the maternal organism, promoting body growth in models that allow this response. Pregnancy-induced body growth appears to be independent of GH action. These findings underscore the need for further studies to investigate the long-lasting consequences of reproductive experience in females.

## Introduction

Pregnancy is a unique state in which numerous physiological adaptations are necessary in the maternal organism to allow proper fetal growth and to prepare for the lactation period (*Augustine et al., 2008*). Among these adaptations, metabolic changes are particularly prominent, including alterations in appetite and glucose homeostasis, as well as increased adiposity (*Augustine et al., 2008*; *Ladyman et al., 2010*; *Zampieri et al., 2015*; *Teixeira et al., 2019a*). Pregnancy also affects neuroplasticity and causes behavioral and emotional alterations (*Larsen and Grattan, 2012*; *Georgescu et al., 2021*). Several of these changes are reversible. However, pregnancy may lead to long-term or permanent consequences. For example, prior pregnancy experience reduces the latency to display maternal behavior in rats (*Bridges, 1978*), indicating long-lasting behavioral adaptations strongly shaped by prior hormonal exposure. Primiparous mice show increased body weight and reduced ambulatory activity compared with age-matched virgin animals, even several months after last contact with the pups (*Ladyman et al., 2018*; *Teixeira et al., 2019b*; *Ladyman et al., 2021*). Body weight retention is frequently observed after pregnancy, especially for women who experienced excessive gestational weight gain (*Scholl et al., 1995*; *Rooney et al., 2005*). Thus, pregnancy may represent a risk factor for obesity in women (*Amorim et al., 2007*). Additionally, reproductive experience induces persistent structural remodeling of the human brain, including reductions in gray matter volume and cortical thickness that remain evident for at least six years after parturition (*Martinez-Garcia et al., 2021*; *Pritschet et al., 2024*). These findings indicate that motherhood is associated with long-term adaptations in both neural and endocrine function (*Bridges, 2016*).

Pregnancy-induced adaptations are thought to be largely driven by the marked rise in circulating hormones during gestation (*Augustine et al., 2008*), including progesterone and estrogens, as well as placentally derived analogues of pituitary hormones, such as chorionic gonadotropin and placental lactogens (also referred to as chorionic somatomammotropins). However, important differences are observed between humans and rodents. Humans, but not rodents, possess the *GH2* gene, which encodes the placental growth hormone (GH) variant, whose production progressively replaces pituitary GH (derived from the *GH1* gene) secretion during pregnancy (*Liao et al., 2018*). In contrast, rodents maintain pituitary GH secretion throughout gestation (*Gatford et al., 2017*; *Liao et al., 2018*). Additionally, human GH, both pituitary and placental variants, as well as chorionic somatomammotropins, can activate both the GH receptor (GHR) and the prolactin receptor (PrlR), explaining their lactogenic effects. In contrast, murine GH activates only the GHR (*Bartke and Kopchick, 2015*).

Despite species-specific differences, it is evident that hormonal exposure during pregnancy not only transiently shapes maternal physiology but may also exert lasting effects on health across the lifespan. However, the chronic consequences of reproductive experience remain poorly understood. Given that most women will experience at least one pregnancy during their lifetime, elucidating how reproductive experience influences long-term physiology and health is of substantial biological and clinical importance. Therefore, the goal of this study is to investigate the long-term effects of reproductive experience on the maternal organism and the potential involvement of the somatotrophic axis in these adaptations.

## Results

### Pregnancy promotes permanent body growth in wild-type and GH-deficient mice

To investigate the consequences of reproductive experience, body weight and composition of C57BL/6J wild-type (WT) mice were initially measured before mating. Pregnant mice gave birth in individual cages and were maintained with their litters for 3 weeks. Two weeks after weaning (the recovery period from lactation), the females were reevaluated to determine changes relative to baseline. The control group consisted of age-matched virgin females. These assessments were also carried out in *Ghrhr^lit/lit^* mice, which carry a null mutation in the gene encoding the GHRH receptor (GHRHR) and are therefore GH-deficient and dwarf. Reproductive experience caused a robust increase in body mass (Figure 1A) and lean mass (Figure 1B) in both control and *Ghrhr^lit/lit^* mice compared with virgin groups. No changes in fat mass were observed between groups (Figure 1C). A new evaluation was performed 2 weeks after the second pregnancy and lactation cycle. Body length measurements revealed a significant increase in multiparous WT and *Ghrhr^lit/lit^* mice compared with age-matched virgin animals (Figure 1D-E). In addition, multiparous mice showed higher body and lean mass than virgin animals (Figure 1F-G). Despite pregnancy-induced growth, *Ghrhr^lit/lit^* mice remained smaller than WT females (Figure 1D-G). Fat mass increased after 2 pregnancies in both WT and *Ghrhr^lit/lit^*mice compared with their respective age-matched virgin groups (Figure 1H). Multiparous WT and *Ghrhr^lit/lit^* mice also had increased liver (Figure 1I) and kidney (data not shown) masses compared with virgin groups. Heart and brain masses increased only in multiparous *Ghrhr^lit/lit^*mice compared with virgin *Ghrhr^lit/lit^* females, whereas these tissues did not change in WT mice (data not shown). Thus, pregnancy promotes body growth in WT and GH-deficient females.

**Figure 1.**
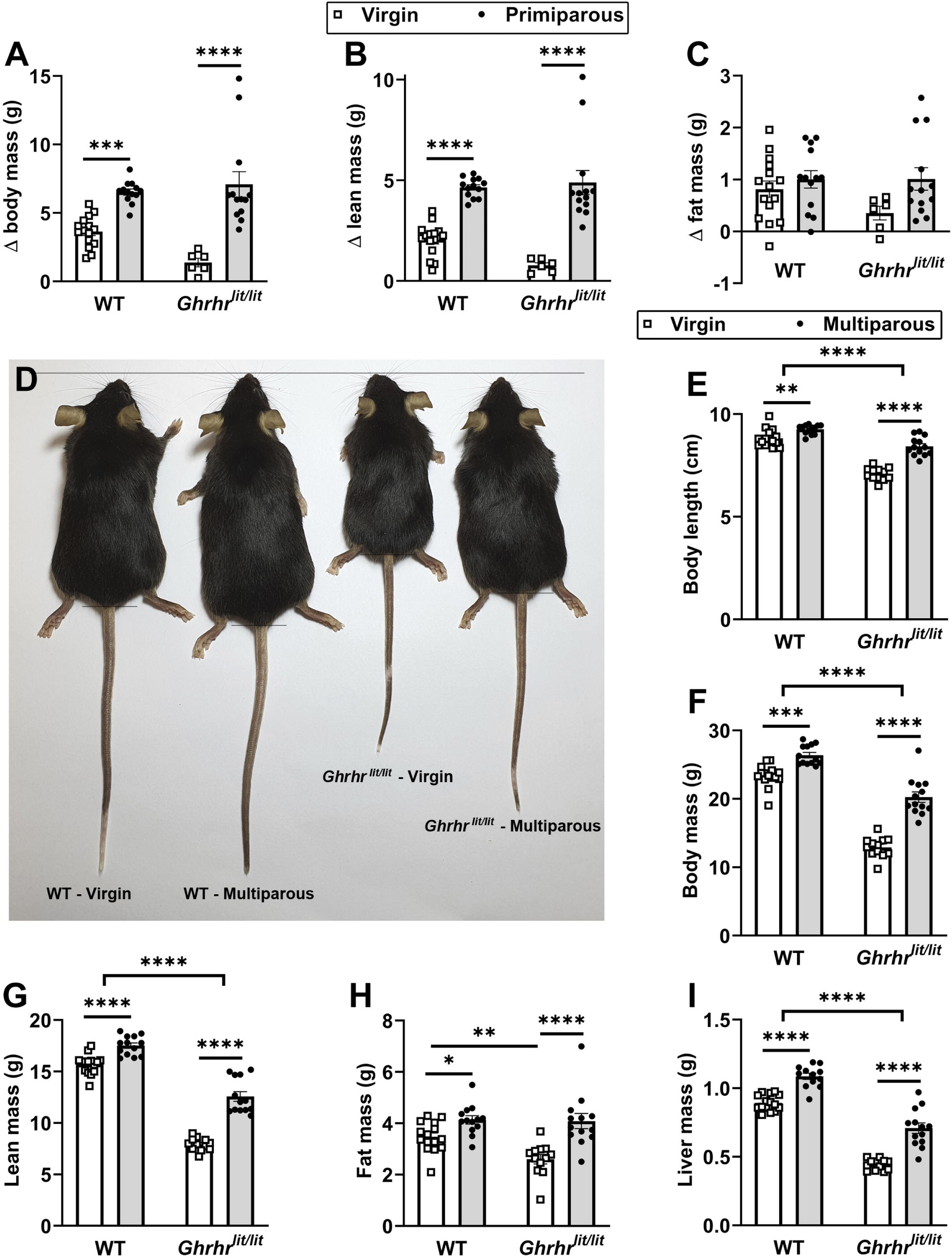
Pregnancy promotes permanent body growth in wild-type and GH-deficient mice. (A-C) Differences relative to baseline in body, lean, and fat mass in virgin WT (n = 15), primiparous wild-type (WT; n = 13), virgin *Ghrhr^lit/lit^*(n = 6), and primiparous *Ghrhr^lit/lit^* (n = 13). (D) Representative image showing differences in longitudinal growth between virgin and multiparous WT and *Ghrhr^lit/lit^* mice. (E-I) Body length, body weight, lean mass, fat mass, and liver mass in multiparous WT (n = 13) and *Ghrhr^lit/lit^* (n = 13) mice, relative to age-matched virgin WT (n = 15) and *Ghrhr^lit/lit^* (n = 12) mice. *, P < 0.05; **, P < 0.01; ***, P < 0.001; ****, P < 0.0001. Statistical analysis was performed using two-way ANOVA and Holm-Sidak’s multiple comparisons test.

### Growth is primarily induced during the first pregnancy

The following experiment tested whether a second pregnancy could also increase parameters indicative of body growth. Unlike the body weight and lean mass gains observed after the first pregnancy, WT mice subjected to a second pregnancy cycle showed no increases in body or lean mass compared with age-matched WT females (Figure 2A-B). Because females undergoing a second pregnancy are inherently older than during their first gestation, we assessed whether a first pregnancy occurring at a comparable age to the second would influence body weight. WT females that had a late first pregnancy (age-matched first pregnancy group) showed increased body and lean mass gain compared with age-matched virgin and second pregnancy groups (Figure 2A-B). Notably, the gains in body and lean mass induced by the first pregnancy were similar among the youngest and oldest females compared with their respective age-matched groups of virgin females (Figure 2A-B). No differences in fat mass gain were observed between the groups (data not shown). In contrast to WT mice, the second pregnancy resulted in a significant increase in body and lean mass in *Ghrhr^lit/lit^* females compared with age-matched virgin animals (Figure 2C-D). However, the magnitude of the increase in the second pregnancy was significantly lower than in the first (Figure 2C-D), highlighting that growth occurs primarily during the first pregnancy.

**Figure 2.**
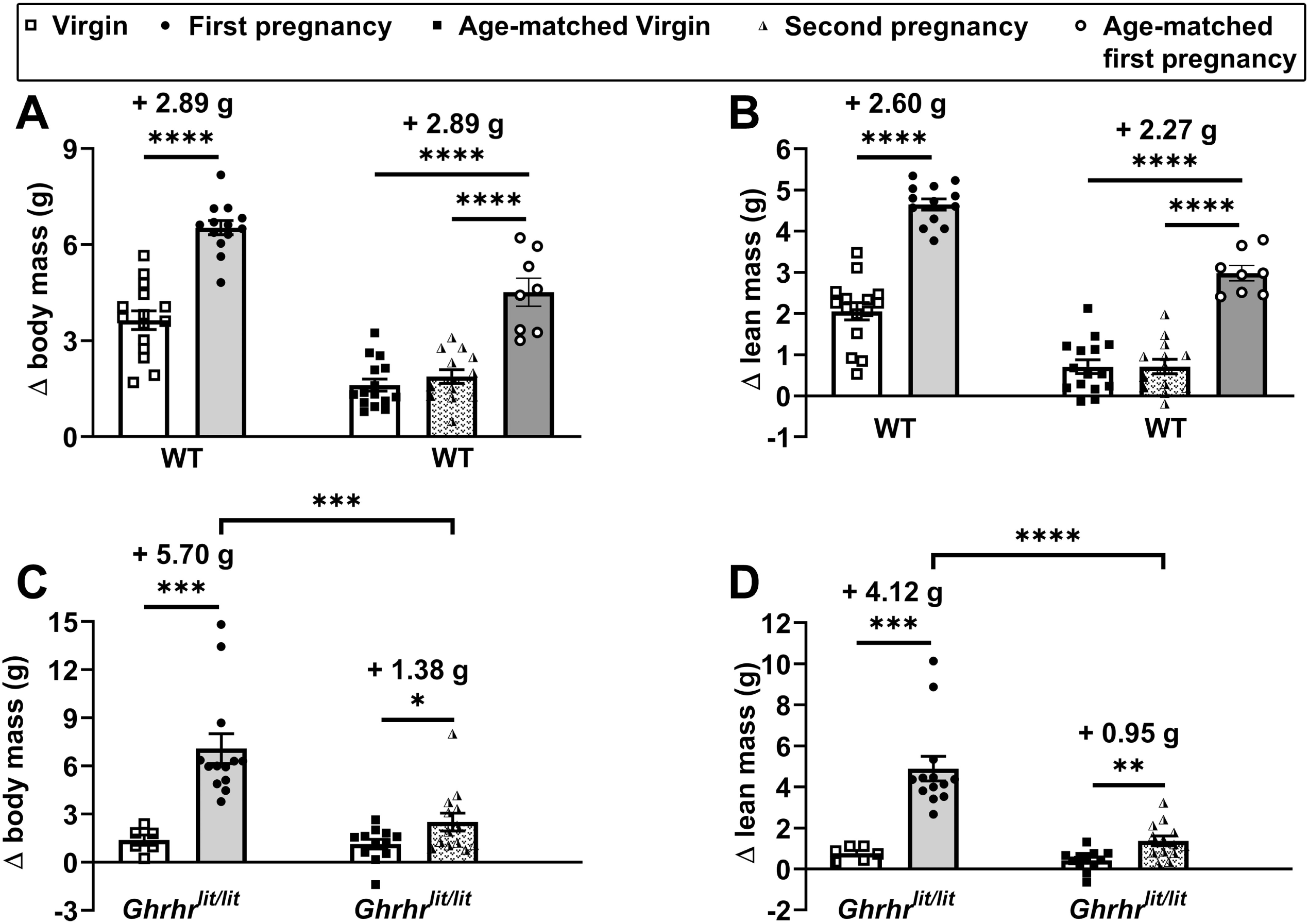
Growth is primarily induced during the first pregnancy. (A-B) Changes in body weight and lean mass in wild-type (WT) virgins (n = 15), after the first pregnancy (n = 13), in virgins matched for age at the second pregnancy (n = 15), after the second pregnancy (n = 13), and after a late first pregnancy (n = 8). (C-D) Changes in body weight and lean mass in *Ghrhr^lit/lit^*virgins (n = 6), after the first pregnancy (n = 13), in virgins matched for age at the second pregnancy (n = 12), and after the second pregnancy (n = 13). *, P < 0.05; **, P < 0.01; ***, P < 0.001; ****, P < 0.0001. Statistical analysis was performed using unpaired two-tailed Student’s t-test or one-way ANOVA and Newman-Keuls multiple comparisons test.

### Increased GH secretion in pregnant WT mice but not in *Ghrhr^lit/lit^*females

Because GH is the major circulating factor that induces longitudinal growth in mammals (*Zhou et al., 1997*; *Dehkhoda et al., 2018*), we investigated whether WT and *Ghrhr^lit/lit^* mice exhibit alterations in GH secretion during pregnancy (gestational age 14-17 days). Confirming previous studies (*Gatford et al., 2017*; *Kaur et al., 2020*; *Wasinski et al., 2022*), pregnancy increased GH secretion in WT mice compared with virgin animals (Figure 3A, C). This increase was associated with higher GH pulse frequency and elevations in both pulsatile and basal secretion (Figure 3C-G). The contribution of basal to total GH secretion increased in pregnant WT mice (Figure 3H). As expected, non-pregnant *Ghrhr^lit/lit^* mice showed blunted GH secretion (Figure 3B). Importantly, pregnancy did not stimulate GH secretion in *Ghrhr^lit/lit^* mice (Figure 3C-H).

**Figure 3.**
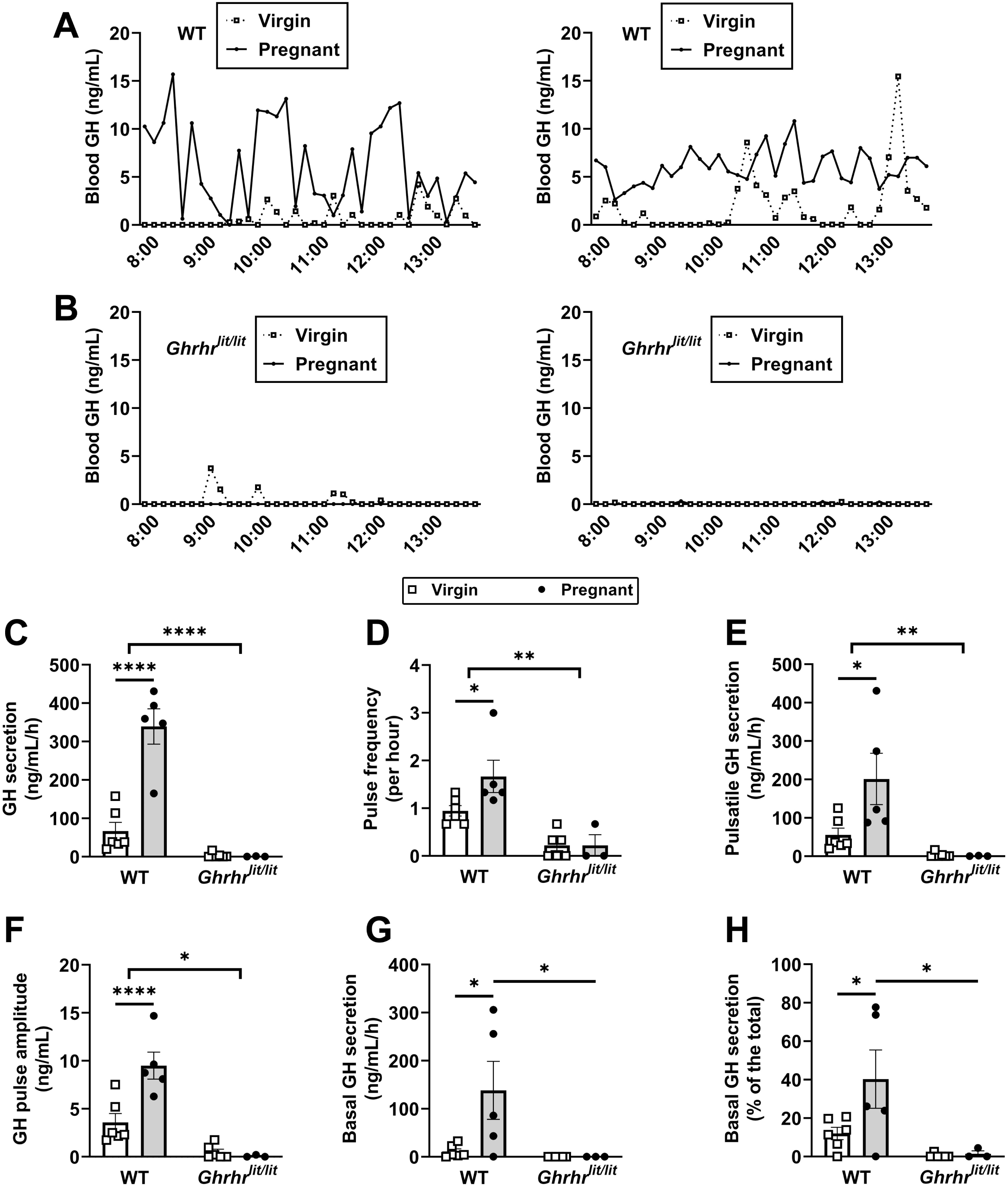
Increased GH secretion in pregnant WT mice but not in pregnant *Ghrhr^lit/lit^* mice. (A-B) Representative examples of the pattern of GH secretion in two pregnant and virgin WT mice and in two pregnant and virgin *Ghrhr^lit/lit^* mice. (C-H). Total GH secretion, GH pulse frequency, pulsatile GH secretion, GH pulse amplitude, basal (non-pulsatile) GH secretion, and contribution of basal secretion to total GH secretion in virgin WT (n = 6), pregnant WT (n = 5), virgin *Ghrhr^lit/lit^* (n = 6), and pregnant *Ghrhr^lit/lit^*(n = 3) mice. The pregnant mice were at gestational ages 14-17 days. *, P < 0.05; **, P < 0.01; ****, P < 0.0001. Statistical analysis was performed using two-way ANOVA and Holm-Sidak’s multiple comparisons test.

### Pregnancy differentially modulates hepatic GHR signaling in WT and *Ghrhr^lit/lit^* mice

GH action in the liver plays a key role in controlling growth because GHR signaling stimulates hepatic insulin-like growth factor 1 (IGF-1) secretion, particularly by activating the signal transducer and activator of transcription 5b (STAT5b) pathway (*Udy et al., 1997*; *Davey et al., 2001*). GH action in the liver is not only controlled by the pattern of GH secretion but also by hepatic GH sensitivity (*Waxman and O’Connor, 2006*; *de Sousa et al., 2025b*). A reduced expression of STAT5b protein was observed in the liver of pregnant WT and *Ghrhr^lit/lit^* mice (Figure 4A,C). Phosphorylation of STAT5 (pSTAT5) was also reduced in the liver of pregnant WT compared to virgin animals (Figure 4B-C). Hepatic pSTAT5 was reduced in *Ghrhr^lit/lit^*mice, compared to virgin WT, and did not change in pregnant animals (Figure 4B-C). Serum IGF-1 and hepatic *Igf1* mRNA levels were reduced in pregnant WT mice compared with virgin animals (Figure 4D-E). In contrast, pregnancy increased serum IGF-1 and hepatic *Igf1* mRNA levels in *Ghrhr^lit/lit^* mice (Figure 4D-E). Corroborating the results of the protein expression, pregnancy reduced *Stat5b* mRNA levels in both WT and *Ghrhr^lit/lit^* mice (Figure 4F). Conversely, hepatic *Ghr* and *Prlr* mRNA levels increased during pregnancy in WT and *Ghrhr^lit/lit^* mice (Figure 4G-H). *Esr1* mRNA expression was reduced in *Ghrhr^lit/lit^* mice and was not affected by pregnancy (Figure 4I). Hepatic *Socs2* and *Cish* RNA levels, transcripts that are indicative of GH action and regulate GH sensitivity (*Greenhalgh and Alexander, 2004*), were also reduced in *Ghrhr^lit/lit^*mice compared to WT animals (Figure 4J-K). Moreover, pregnancy increased *Socs2* and *Cish* RNA expression only in *Ghrhr^lit/lit^*mice (Figure 4J-K). Altogether, although WT mice exhibit increased GH secretion and hepatic *Ghr* expression, molecular analyses do not indicate higher GH action in the liver. In contrast, *Ghrhr^lit/lit^* mice showed evidence of greater GH action in the liver, including increased circulating and hepatic IGF-1 levels, as well as greater *Socs2* and *Cish* expression.

**Figure 4.**
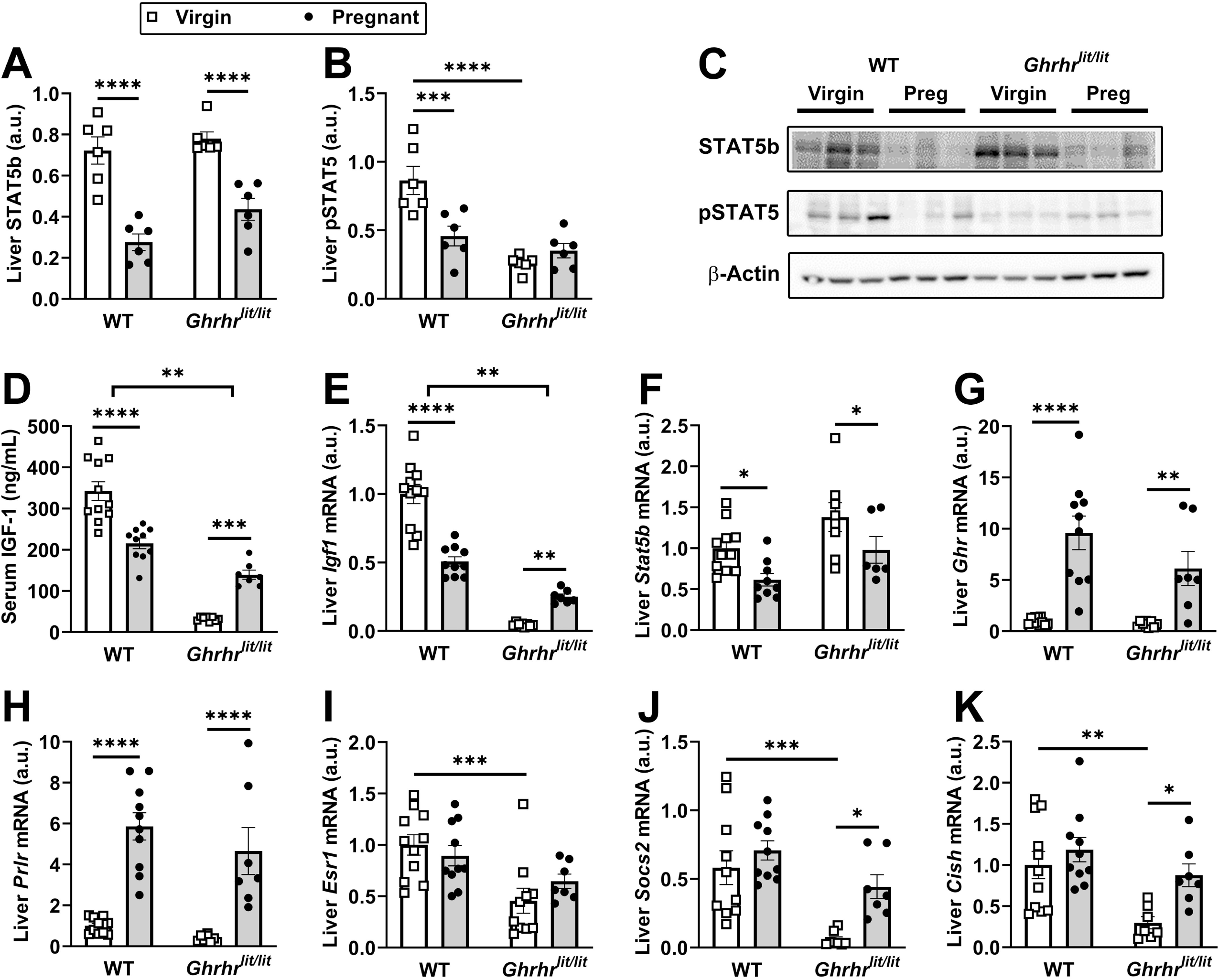
Pregnancy differentially modulates hepatic GHR signaling in WT and *Ghrhr^lit/lit^* mice. (A-C) Western blot analysis to determine the hepatic expression of STAT5b and pSTAT5 proteins in virgin WT (n = 6), pregnant WT (n = 6), virgin *Ghrhr^lit/lit^* (n = 6), and pregnant *Ghrhr^lit/lit^* (n = 6) mice. (D-K) Serum IGF-1 levels and hepatic mRNA expression in virgin WT (n = 11), pregnant WT (n = 10), virgin *Ghrhr^lit/lit^* (n = 8), and pregnant *Ghrhr^lit/lit^*(n = 7) mice. The pregnant mice were at gestational ages 14-17 days. *, P < 0.05; **, P < 0.01; ***, P < 0.001; ****, P < 0.0001. Statistical analysis was performed using two-way ANOVA and Holm-Sidak’s multiple comparisons test.

### Pregnancy-induced body growth is preserved despite disruption of GH-, ghrelin-, and estrogen-related signaling pathways

To gain further insights into the mechanisms driving post-pregnancy body growth, the effects of pregnancy were investigated across different mouse models. Mice carrying a GHR deletion in hepatocytes (Alb^ΔGHR^) show blunted circulating IGF-1 levels and growth deficits despite higher GH secretion (*List et al., 2014*; *Wasinski et al., 2023*; *de Sousa et al., 2025b*). We observed that a first pregnancy caused a robust increase in body weight and lean mass in Alb^ΔGHR^ mice, which was comparable to that seen in control animals (Figure 5A-B). Alb^ΔGHR^ females also showed increased body weight after the second pregnancy, but this was explained by pregnancy-induced increases in fat mass (Figure 5C). Thus, GHR signaling in hepatocytes is not necessary to induce pregnancy-induced growth. Ghrelin is a GH secretagogue, and impaired ghrelin signaling may affect body growth (*Peris-Sampedro et al., 2021*; *Punt et al., 2025*). However, ghrelin receptor-deficient mice (GHSR KO) showed increases in body weight and lean mass similar to those of control animals (Figure 5D-E). Sex hormones also regulate growth, and estrogen receptor α (ERα) signaling activates the STAT5 pathway in the liver and IGF-1 synthesis (*Venken et al., 2005*). In addition, estradiol treatment stimulates growth in GHR-deficient mice (*Venken et al., 2005*), and estrogen response elements are found in the mouse *Stat5b* gene (*Furigo et al., 2014*). Nonetheless, hepatocyte-specific ERα ablation did not prevent pregnancy-induced increases in body weight, lean mass, and adiposity (Figure 5F-H). We also evaluated mice carrying brain GHR inactivation, as this is a model of gigantism caused by loss of GH negative feedback (*Furigo et al., 2019*; *Teixeira et al., 2019a*; *Wasinski et al., 2020*; *de Sousa et al., 2025a*). First pregnancy increased body, lean, and fat mass in Nestin^ΔGHR^ females, as observed in control animals (Figure 5I-K).

**Figure 5.**
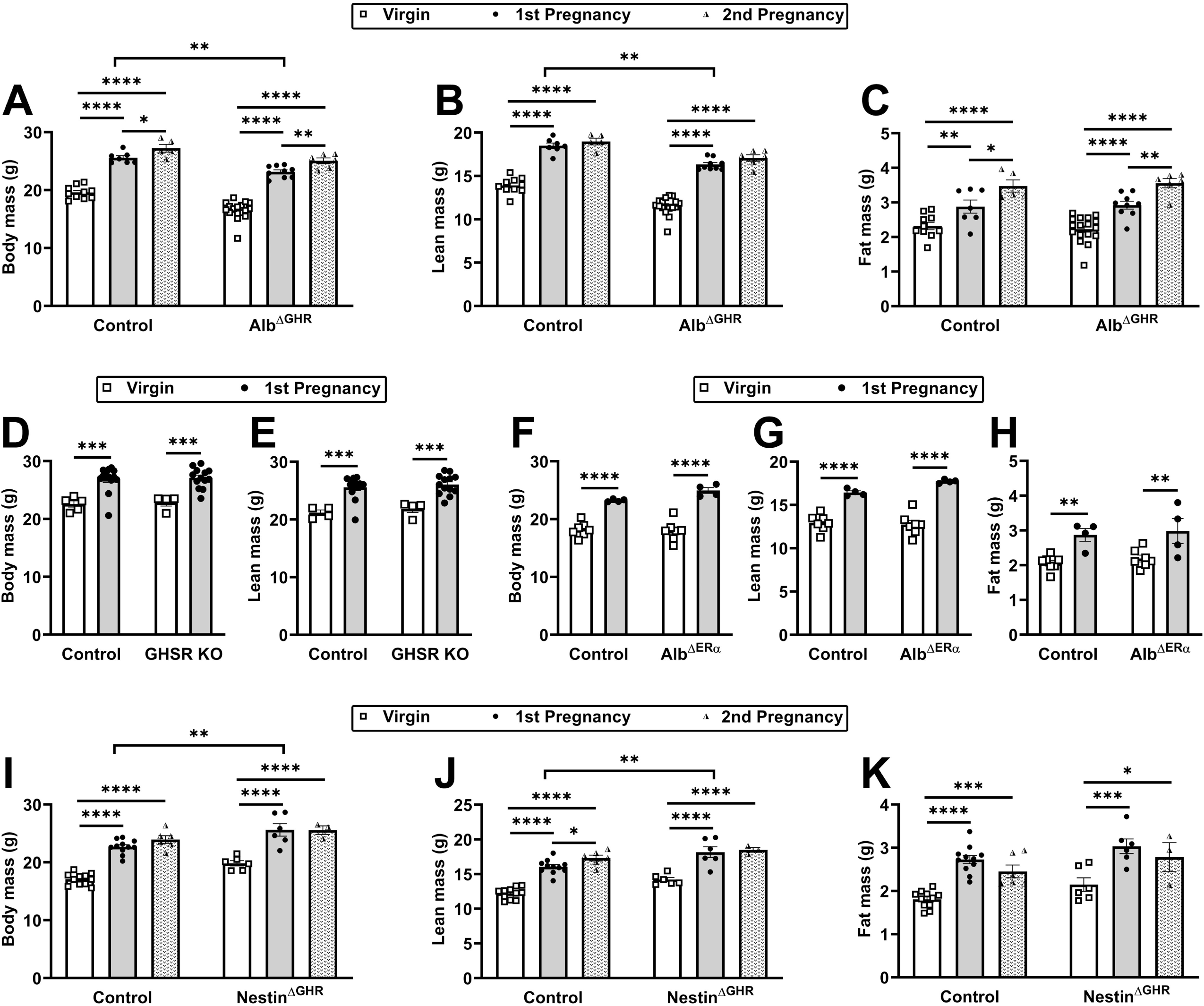
Pregnancy-induced body growth is preserved despite disruption of GH-, ghrelin-, and estrogen-related signaling pathways. (A-C) The effects of first and second pregnancies on body weight, lean mass, and fat mass in control virgin (n = 10), control 1^st^ pregnancy (n = 7), control second pregnancy (n = 5), Alb^ΔGHR^ virgin (n = 17), Alb^ΔGHR^ 1^st^ pregnancy (n = 9), and Alb^ΔGHR^ second pregnancy (n = 6). (D-E) Body weight and lean mass in control virgin (n = 5), control 1^st^ pregnancy (n = 14), GHSR KO virgin (n = 4), and GHSR KO 1^st^ pregnancy (n = 13). (F-H) Body weight, lean mass, and fat mass in control virgin (n = 7), control 1^st^ pregnancy (n = 4), Alb^ΔERα^ virgin (n = 7), and Alb^ΔERα^ 1^st^ pregnancy (n = 4). (I-K) The effects of first and second pregnancies on body weight, lean mass, and fat mass in control virgin (n = 11), control 1^st^ pregnancy (n = 11), control second pregnancy (n = 6), Nestin^ΔGHR^ virgin (n = 6), Nestin^ΔGHR^ 1^st^ pregnancy (n = 6), and Nestin^ΔGHR^ second pregnancy (n = 3). *, P < 0.05; **, P < 0.01; ***, P < 0.001; ****, P < 0.0001. Statistical analysis was performed using two-way ANOVA and Holm-Sidak’s multiple comparisons test.

### Case reports of Itabaianinha women with isolated GH deficiency suggest post-pregnancy growth

Growth plate physiology differs between humans and rodents: growth plates remain open after sexual maturation in rodents, whereas they fuse at the end of puberty in humans, thereby terminating longitudinal bone growth (*Emons et al., 2011*). Although pregnancy-induced longitudinal growth is not observed in normal adult women, we speculate that a slight pregnancy-induced growth may happen in certain special cases. To provide proof-of-concept evidence that gestational factors permanently influence the maternal organism in humans, we studied a cohort of individuals with severe isolated GH deficiency (IGHD) caused by a homozygous loss-of-function mutation (c.57+1G→A) in the GHRHR gene. These individuals are from the Brazilian city of Itabaianinha (*Salvatori et al., 1999*; *Aguiar-Oliveira and Salvatori, 2021*), and their mutation replicates effects seen in *Ghrhr^lit/lit^* mice (*Aguiar-Oliveira and Bartke, 2019*).

Using data from this rare cohort, we compared the height of 8 nulliparous women and 9 subjects with one or more pregnancies, both groups with IGHD without lifetime GH replacement therapy. The average number of children per woman was 1.8 children (*Aguiar-Oliveira and Salvatori, 2021*). Nulliparous women were 7 cm shorter than those with one or more pregnancies, 1.13 ± 0.04 m vs 1.20 ± 0.05 m (P = 0.009; Figure 6A). Moreover, the three shortest women, with heights of 1.07, 1.08, and 1.11 m, were nulliparous. Anecdotal reports from 6 IGHD cases also suggest residual growth in 4 women after pregnancy (Table 1). This information was obtained through face-to-face personal interviews with one of the researchers (M.H. A.-O.), who was also responsible for measuring height before and after pregnancy. Subject 1 reported clear and consistent signs of somatic growth following two pregnancies. She is married to an IGHD individual, and both of her sons (currently 30 and 33 years old) are affected, as expected for this genetic background. According to documented measurements, her height increased from 1.16 m before pregnancy to 1.20 m after her two pregnancies. She also reported an increase in shoe size from 25 to 28 (Brazilian size) and changes in ring size, as a ring previously worn on the fourth finger no longer fit after childbirth. Together, these findings are consistent with clinically perceptible post-pregnancy somatic growth.

**Figure 6.**
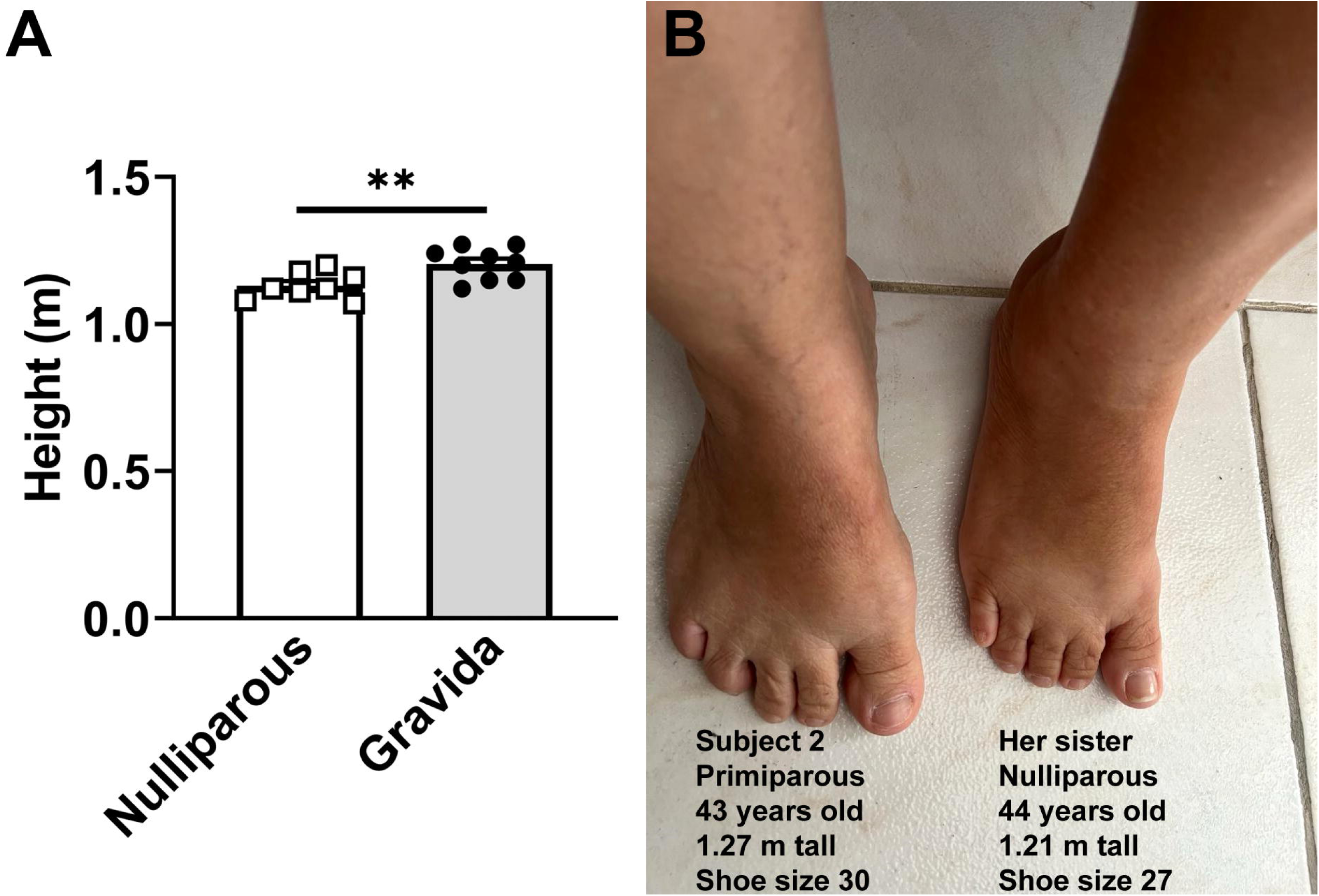
Evidence of post-pregnancy growth in women with isolated GH deficiency. (A) In a cohort of women with isolated GH deficiency caused by a loss-of-function mutation in the GHRHR gene, nulliparous women (n = 8) were shorter than those with one or more pregnancies (n = 9). **, P = 0.009 (unpaired two-tailed Student’s t-test). (B) Comparison of foot size between two sisters with IGHD, one nulliparous and another primiparous. Image of the feet of two sisters with severe isolated GH deficiency caused by a homozygous loss-of-function mutation in the GHRHR gene. In addition to a larger shoe size, subject 2, who had a heterozygous son 9 years ago, is also taller than her nulliparous sister.

**Table 1.** Characteristics of the six subjects with IGHD who had one or more pregnancies.

|  | Age at the time of pregnancy | Height before pregnancy | Height after pregnancy |
| --- | --- | --- | --- |
| Subject 1 | 27 years old<sup>2</sup> | 1.16 m | 1.20 m |
| Subject 2 | 34 years old<sup>1</sup> | 1.26 m | 1.27 m |
| Subject 3 | 23 years old<sup>1</sup> | 1.44 m | 1.47 m |
| Subject 4 | 25 years old<sup>1</sup> | 1.40 m | 1.42 m |
| Subject 5 | 27 years old<sup>3</sup> | N/A | 1.24 m |
| Subject 6 | 24 years old<sup>2</sup> | N/A | 1.27 m |

<sup>1</sup> first pregnancy; <sup>2</sup> second pregnancy; <sup>3</sup> third pregnancy; N/A not available.

Three additional women with IGHD also reported body changes suggestive of growth after pregnancy, though less noticeable than those of subject 1. Subject 2 (43 years old) had a documented increase in height from 1.26 m before pregnancy to 1.27 m at the present. She has one heterozygous son (8.8 years old). She reported an increase in shoe size from 29 to 30, and her graduation ring no longer fits; however, she also gained weight, which may have contributed to the ring size changes. Her feet are larger than her sister’s; she is 44 years old, 1.21 m tall, shoe size 27, and nulliparous (Figure 6B). Subject 3, treated with GH in childhood and early adolescence, was 1.44 m tall before pregnancy. She had a heterozygous boy 13 years ago; now she is 36 years old and 1.47 m tall. Subject 4, currently 28 years old, was treated with GH in childhood and early adolescence. She had a heterozygous girl 3 years ago. Her current height is 1.42 m, but before pregnancy, she was 1.40 m tall. In contrast, 2 sisters with IGHD reported no signs of pregnancy-induced growth. Subject 5, 1.24 m tall, aged 56, has a daughter, aged 35, and two sons, aged 30 and 29, each heterozygous. Subject 6, 1.27 m tall, aged 68, has two heterozygous daughters, aged 46 and 44, respectively. However, no information about their heights before pregnancy is available. Despite these two cases, our data indicate a slight post-pregnancy growth in women with severe IGHD.

## Discussion

Pregnancy causes profound endocrine and metabolic changes that are conserved across mammals. The maternal organism undergoes significant hormonal shifts that not only temporarily alter the body but may also lead to lasting effects. Here, we demonstrated that reproductive experience resulted in increased body growth in WT mice, independent of GH. This growth was mainly observed after the first pregnancy, while subsequent reproductive experiences primarily increased body fat. In humans, puberty triggers the fusion of growth plates, leading to the end of longitudinal bone growth (*Emons et al., 2011*). Height gain is typically limited after menarche, averaging 6.6-8.94 cm depending on the studied cohort (*Spear, 2002*; *Gardstedt-Berghog et al., 2024*; *Gaete et al., 2025*; *Xu et al., 2026*). Previous studies indicate that pregnant women may experience a slight but significant decrease in height due to weight gain, vertebral compression, and increased lordosis (*Scholl et al., 1993*; *Xu et al., 2026*). Therefore, using height as a measure of post-pregnancy growth in humans may be unreliable, which may explain why normal gravidas do not show longitudinal growth. However, increased foot size has been reported after pregnancy (*Wetz et al., 2006*; *Dunn et al., 2012*). Despite the lack of evidence for post-pregnancy longitudinal growth in healthy women, the current results in mice suggest that pregnancy activates GH-independent growth pathways. In species with open growth plates, like rodents, this could result in actual longitudinal growth, whereas in humans, it may instead lead to tissue remodeling or physical changes that do not increase height but may have lasting effects that are still not well understood. Thus, the mouse model offers mechanistic insight into conserved pregnancy-related anabolic processes that may operate independently of the somatotropic axis.

Multiparous *Ghrhr^lit/lit^* mice showed significant gains in weight (+57%) and lean mass (+59%) after pregnancy compared with age-matched *Ghrhr^lit/lit^* virgins. In fact, their relative weight and lean mass gains were much higher than those observed in WT mice, which showed ∼13% more body weight and lean mass than WT virgins. Thus, GH-deficient mice are considerably more sensitive to pregnancy-induced growth than WT mice. The profound growth deficit in *Ghrhr^lit/lit^* mice likely enhances their sensitivity to pregnancy-induced growth factors. Human data are in accordance with these findings as children with severe GH deficiency (GHD) show a higher response to GH replacement than patients with less severe or questionable GHD (*Savage and Bang, 2012*; *Wit et al., 2019*). Accordingly, patients exhibiting the lowest response during GH provocation tests showed the highest responses to GH replacement treatment (*Donbaloglu et al., 2023*). Regarding the capacity of women with IGHD to present post-pregnancy growth, residual spinal growth is frequently observed after skeletal maturity, particularly in the 1–2 years following the peak of puberty, often averaging just a few millimeters to 1 cm in height (*Ghanem and Rizkallah, 2020*). The increased responsiveness to growth factors, as well as the late menarche (±17 years) and, therefore, delayed bone maturation (*Aguiar-Oliveira and Salvatori, 2021*), can also explain why adult women with severe IGHD retained some growth capacity after pregnancy. Importantly, identifying adult women with untreated IGHD who subsequently experienced pregnancy is extremely challenging. GH replacement therapy is typically initiated early in life, and untreated individuals are rare. Therefore, although a limited number of cases were available for study and the human data relied in part on participants’ self-reported information, this sample represents a highly valuable and unique clinical resource.

Unlike WT females, *Ghrhr^lit/lit^* mice show no increase in GH secretion during pregnancy, maintaining suppressed levels similar to those of nulliparous *Ghrhr^lit/lit^* mice. Thus, these findings revealed that GHRH signaling seems required for pregnancy-induced increases in GH secretion in mice. However, we cannot distinguish whether the lack of pregnancy-induced increases in GH secretion reflects the absence of GHRH signaling per se or developmental defects resulting from GHRH deficiency. More importantly, this result indicates that pregnancy-induced body growth is independent of GH levels. It is important to note that, despite marked species-specific differences in the GH locus (as summarized in the Introduction), pregnancy-induced growth appears to have been evolutionarily conserved. Although this response may be mediated by distinct mechanisms in rodents and humans, the available evidence suggests that it is largely GH-independent in both species.

However, the expression of genes typically induced by GHR signaling, such as *Igf1*, *Socs2*, and *Cish*, was upregulated in the liver of pregnant *Ghrhr^lit/lit^* mice compared with virgin animals. The increased circulating IGF-1 levels in *Ghrhr^lit/lit^*mice may have contributed to their greater relative growth during gestation. In contrast, pregnant WT mice showed decreased circulating IGF-1 levels and hepatic *Igf1* mRNA expression compared to virgin animals, suggesting hepatic GH resistance. Indeed, hepatic STAT5/GH signaling desensitization was described in pregnant mice, which involves increased expression of negative regulators of STAT signaling (*Miquet et al., 2005*), as observed in the current study. These results are consistent with another study that also observed decreased IGF-1 secretion and hepatic *Igf1* mRNA expression in mid-pregnant mice (*Travers et al., 1990*). In humans, IGF-1 levels decrease during the first trimester of gestation (*Caron et al., 2010*; *Persechini et al., 2015*), but increase during mid- and late-pregnancy, when they correlate with placental GH levels (*Caufriez et al., 1993*). Altogether, pregnancy differentially modulates the hepatic expression of genes commonly induced by GHR signaling in WT and *Ghrhr^lit/lit^* mice. Thus, pregnancy-induced growth may occur through distinct mechanisms in WT and *Ghrhr^lit/lit^*mice. Whereas increased circulating IGF-1 levels may contribute to growth in *Ghrhr^lit/lit^* mice, pregnancy-induced growth in WT mice may be mediated through liver-derived IGF-1-independent mechanisms. Locally produced IGF-1 in tissues such as the growth plate, bone, and skeletal muscle may be regulated by gestational factors, thereby promoting body growth. Future studies are needed to investigate this possibility.

In an attempt to identify specific hormonal factors involved in pregnancy-induced body growth, mice with hepatocyte-specific deletion of GHR or ERα were generated. Confirming previous findings that pregnancy-induced body growth is independent of GH, hepatic GHR deletion did not prevent increases in body weight and lean mass in primiparous and multiparous mice. Although estradiol treatment can stimulate growth in GHR-deficient mice and activate the STAT5 pathway and IGF-1 synthesis in the liver (*Venken et al., 2005*), hepatocyte-specific ERα KO mice also exhibited normal pregnancy-induced body growth. Similar findings were observed in GHSR-deficient mice, demonstrating that pregnancy-induced body growth is ghrelin-independent. Finally, a gigantism model (Nestin^ΔGHR^ mice) was tested to determine whether pregnancy-induced growth can occur in mice that already have increased baseline lean body mass (+16%) compared with control animals. However, pregnancy also increased body and lean mass in Nestin^ΔGHR^ mice, in a manner comparable to control animals, suggesting that this growth potential persists even in animals that begin gestation with greater somatic length.

Several questions arose from our findings. For example, why does body growth occur primarily after the first pregnancy, regardless of the age at which it occurs? We speculate that pregnancy hormones, particularly sex hormones, may modify the growth plate of long bones, limiting further growth in subsequent gestations. Future studies should investigate the impact of pregnancy on growth plate structure and function. Another interesting finding is that multiple gestations are associated with increased body adiposity in mice. Leptin resistance is a hallmark of pregnancy, favoring physiological increases in food intake and fat mass (*Mounzih et al., 1998*; *Trujillo et al., 2011*; *Ladyman et al., 2012*; *Zampieri et al., 2015*; *Andreoli et al., 2019*; *Gustafson et al., 2019*). If leptin resistance persists in the postpartum period, it may favor fat retention. However, previous studies indicate normal leptin sensitivity in female mice following pregnancy (*Ladyman et al., 2018*; *Andreoli et al., 2019*; *Teixeira et al., 2019b*). It is also possible that each pregnancy may cause some degree of fat retention, so multiple pregnancies lead to a significant increase in body adiposity simply through an additive effect. Regardless of the mechanism that causes pregnancy-induced fat gain, it is evident that pregnancy may represent a risk factor for obesity (*Scholl et al., 1995*; *Rooney et al., 2005*; *Amorim et al., 2007*).

Our study was not intended to distinguish the effects of pregnancy from those of lactation. Lactation stimulates the secretion of prolactin and oxytocin (*Augustine et al., 2008*; *Ladyman et al., 2010*). In mice, lactation is a period of high metabolic demand due to the cost of milk production, which is compensated for by hyperphagia (*Woodside et al., 2012*; *Zampieri et al., 2015*; *Teixeira et al., 2019a*). Thus, lactation may have contributed to the growth of the females and should be considered alongside gestation when interpreting our results. Moreover, female mice housed with sterile males exhibit increased body weight and reduced lifespan (*Garratt et al., 2020*). Therefore, mating per se, even in the absence of fertilization, is sufficient to influence body weight and longevity, representing an additional factor—beyond pregnancy and lactation—that may have contributed to the parameters analyzed.

Since circulating GH and IGF-1 levels do not fully explain pregnancy-induced growth, several additional endocrine factors may contribute to this phenomenon, each potentially accounting for part of the observed response. Prolactin and placental lactogens (chorionic somatomammotropins in humans) are particularly attractive candidates because they activate intracellular signaling pathways that overlap with those of GH, stimulate cell proliferation, play central roles in maternal physiological adaptations, are markedly elevated during pregnancy, and act on multiple target tissues (*Augustine et al., 2008*; *Ladyman et al., 2010*). Therefore, these hormones may contribute to the increased body growth observed in reproductively experienced females. IGF-2 represents another potential mediator. Although best known for its critical role in fetal growth, IGF-2 is abundantly produced at the maternal-fetal interface (*Lopez-Tello et al., 2023*). While its role in pregnancy-induced maternal growth remains unknown, mice with persistent postnatal IGF-2 expression exhibit increased body growth despite normal circulating IGF-1 concentrations (*Younis et al., 2026*). Moreover, IGF-2 contributes to maternal adaptations during pregnancy, including the development of mild insulin resistance (*Lopez-Tello et al., 2023*), making it an attractive candidate to mediate pregnancy-induced body growth. Finally, insulin is a well-established anabolic hormone whose secretion increases during pregnancy (*Augustine et al., 2008*; *Ladyman et al., 2010*). Thus, enhanced insulin receptor signaling may also directly or indirectly contribute to pregnancy-induced maternal growth. Investigating the contribution of these factors to the long-term consequences of reproductive experience, particularly the body growth observed in the present study, represents an important direction for future research.

In conclusion, reproductive experience, including mating, pregnancy, and lactation, activates redundant, multi-hormonal anabolic programs involving growth factors and placental lactogens, among others. Our findings indicate that this coordinated endocrine-metabolic environment is sufficient to promote longitudinal skeletal growth, even in the absence of pituitary GH, in species with open growth plates. We also provided evidence that post-pregnancy growth can occur in women with severe IGHD. Thus, reproductive experience has many effects on the maternal organism. Despite the lack of longitudinal growth in normal women, their bodies undergo significant and potentially permanent changes that need further study to understand the long-term impact on women’s health.

## Material and Methods

### Animals

In the current study, the following mouse strains were used: C57BL/6J (The Jackson Laboratory, Bar Harbor, ME; RRID: IMSR_JAX:000664), *Ghrhr^lit/lit^*(The Jackson Laboratory; RRID: IMSR_JAX:000533), Alb^Cre^ (The Jackson Laboratory; RRID: IMSR_JAX:003574), Ghr^flox^ (*List et al., 2014*), ERα^flox^ (The Jackson Laboratory; RRID: IMSR_JAX:032173), Nestin^cre^ (The Jackson Laboratory; RRID: IMSR_JAX:003771), and GHSR-deficient mice (*Zigman et al., 2005*). The generation, genotyping, and validation of the experimental mice were carried out as described in previous studies from our group (*Frazao et al., 2013*; *Furigo et al., 2019*; *Teixeira et al., 2019a*; *Wasinski et al., 2020*; *Wasinski et al., 2022*; *Wasinski et al., 2023*; *de Sousa et al., 2025a*; *de Sousa et al., 2025b*). Mice had ad libitum access to regular rodent chow (2.99 kcal/g, 9.4% kcal from fat, 236 g protein/kg; Nuvilab CR-1, Quimtia, Brazil) and filtered water. The experimental procedures were approved by the Ethics Committee on the Use of Animals of the Institute of Biomedical Sciences at the University of São Paulo.

### Evaluation of pregnancy-induced growth in mice

The body composition (lean and fat mass) of adult (∼10-week-old) female mice was determined by time-domain nuclear magnetic resonance using the LF50 body composition mice analyzer (Bruker, Germany). Subsequently, the females were mated with sexually experienced males. Upon detection of pregnancy, females were placed in individual cages and maintained with their litters for 3-4 weeks. After weaning, the dams were regrouped in cages with up to five females. Body composition was reassessed 2 weeks after the last contact with the pups (∼20-week-old), which we consider the lactation recovery period. At the same time, body composition was analyzed in age-matched virgin females as a control group. In some experiments, the primiparous females were allowed to mate again, and the entire procedure was repeated to determine the effects of the second pregnancy and lactation cycle. After the last body composition assessment, WT and *Ghrhr^lit/lit^* mice were euthanized in a CO_2_ chamber, and naso-anal length was measured, as were the masses of the liver, heart, kidneys, and brain. Figure 7 summarizes the experimental timeline. Specifically, for GHSR KO mice and their respective controls, lean mass was calculated as total body weight minus fat mass (the sum of periuterine, periovarian, retroperitoneal, inguinal, posterior subcutaneous, and anterior subcutaneous fat depots).

**Figure 7.**
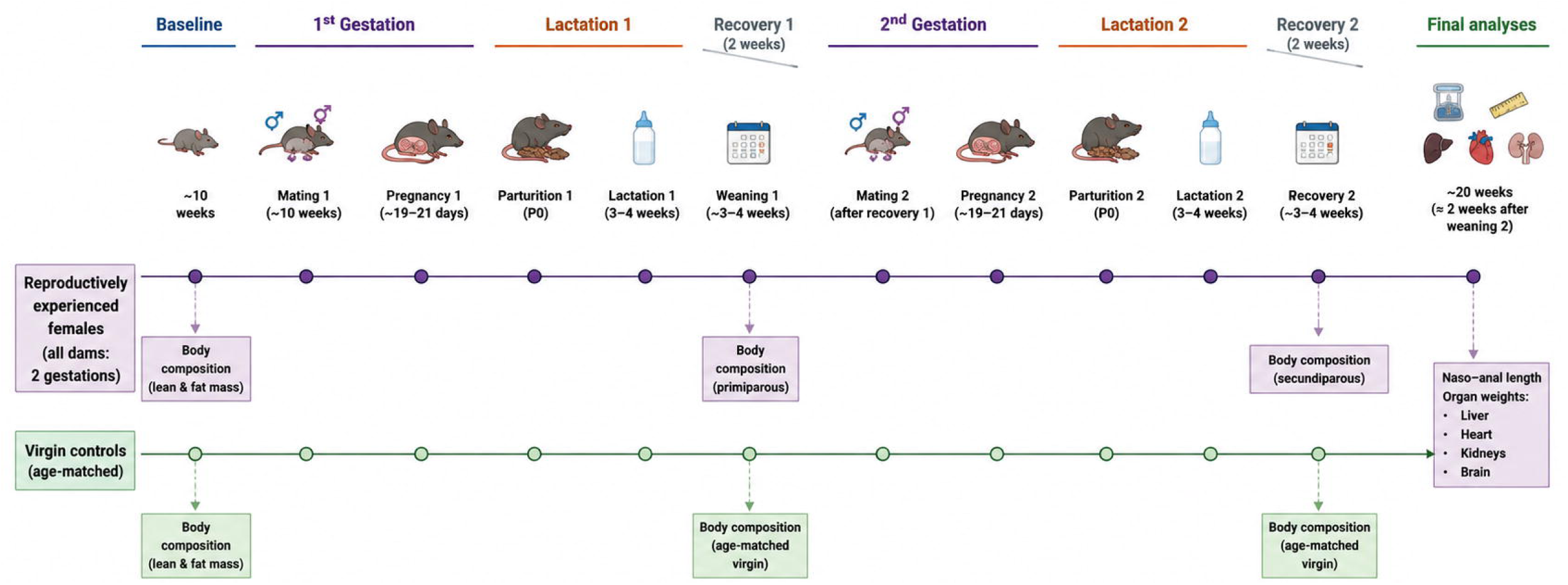
Experimental timeline illustrating the study design. The schematic was created with the assistance of OpenAI image-generation tools and subsequently reviewed and edited by the authors for scientific accuracy.

### Evaluation of pulsatile GH secretion and IGF-1 levels

To adapt the mice to the tail-tip blood sampling procedure, females were acclimated daily for 2-3 weeks before being placed in breeding. This adaptation continued while the females were housed with males. In pregnant females (gestational age 14-17 days) and virgin controls, a total of 36 blood samples were taken from the tail tip every 10 minutes. The procedure initially involved clipping a small portion (1 mm) from the tail tip with a surgical blade. Each blood sample (5 μL for WT and 2 μL for *Ghrhr^lit/lit^*) was then transferred to a tube containing PBS with 0.05% Tween-20. After each draw, gentle fingertip pressure was applied to the tail to halt bleeding. The mice continued to move freely within their home cages, with unlimited access to food and water. Immediately after collection, samples were placed on dry ice and stored at -80°C. An enzyme-linked immunosorbent assay (ELISA) was performed to measure blood GH levels, using a 1:22 final dilution, and the following antisera: monkey anti-rat GH as the capture antibody (1:50,000; NHPP, Cat# AFP411S; RRID: AB_2665564) and rabbit anti-rat GH as the detection antibody (1:100,000; NHPP, Cat# AFP5672099, RRID: AB_2721132). GH pulses were detected using the DynPeak pulse detection algorithm’s default settings (*Vidal et al., 2012*). More details about this ELISA and the analysis of the GH secretion pattern are found in previous publications (*Wasinski et al., 2020*; *Wasinski et al., 2022*; *de Sousa et al., 2024*; *de Sousa et al., 2025b*). Serum IGF-1 levels were measured with a commercial ELISA kit (#MG100; RRID: AB_2827989; R&D Systems, Minneapolis, MN, USA).

### Western Blot

Liver proteins were extracted using RIPA buffer (Sigma-Aldrich, Cat# R0278), containing phosphatase (Sigma-Aldrich, Cat# P5726 and P0044) and protease (Sigma-Aldrich, Cat# P8340) inhibitors. Samples were fractionated by polyacrylamide gel electrophoresis containing SDS (SDS-PAGE) using a running buffer (25 mM Tris; 192 mM Glycine; 0.1% SDS). Then, proteins were transferred to a nitrocellulose membrane overnight at 15 V using transfer buffer (25 mM Tris; 192 mM Glycine; 20% Methanol). Membranes were blocked at room temperature for 2 h in 5% skim milk diluted in TBS-T buffer (10 mM Tris–HCl; 150 mM NaCl; 0.05% Tween-20). Membranes were incubated overnight in primary antibodies diluted in TBS-T containing 3% albumin. Membranes were washed three times for 10 min with TBS-T and incubated at room temperature for 1 h with respective secondary antibodies, diluted in 5% skim milk in TBS-T. After three 10-minute washes with TBS-T, the membranes were treated with chemiluminescent reagents (Clarity Western ECL Substrate, Bio-Rad, Hercules, CA; Cat# 170-5060). Bands on the membranes were revealed by automatic imaging systems. The intensities of the bands were quantified by densitometry using ImageJ software (https://imagej.net/ij/) and normalized to total protein expression in the gel (Ponceau staining). Primary antibodies used for blotting were anti-phospho^Tyr694^-STAT5 antibody (Cell Signaling Technology, Beverly, MA; Cat# 9351; RRID: AB_2315225; 1:1,000), anti-STAT5b antibody (Santa Cruz, Dallas, TX; Cat# sc-377069; 1:1,000), and anti-β-actin (Santa Cruz; Cat# sc-47778, RRID: AB_626632; 1:2,500).

### Gene expression

Total RNA from the liver was extracted with TRIzol (Invitrogen, Carlsbad, CA), followed by incubation in DNase I RNase-free (MilliporeSigma, St. Louis, MO, USA). Reverse transcription was performed using 2 µg of total RNA, SuperScript II Reverse Transcriptase (Invitrogen), and random primers p(dN)6 (MilliporeSigma). Real-time PCR was performed using the 7500TM Real-Time PCR System (Applied Biosystems, Warrington, UK) and SYBR Green Gene Expression PCR Master Mix (Applied Biosystems. The following primers were used: *Actb* (forward: gctccggcatgtgcaaag; reverse: catcacaccctggtgccta), *Cish* (forward: tcgggaatctgggtggtact; reverse: taggaatgtaccctccggca), *Esr1* (forward: gcagatagggagctggttca; reverse: tggagattcaagtccccaaa), *Ghr* (forward: atcaatccaagcctggggac; reverse: acagctgaatagatcctgggg), *Igf1* (forward: ccacactgacatgcccaaga; reverse: gtacttcctttccttctcctttgc), *Prlr* (forward: cagtaaatgccaccaacgaa; reverse: gaggaggctctggttcaaca), *Ppia* (forward: tatctgcactgccaagactgagt; reverse: cttcttgctggtcttgccattcc), *Socs2* (forward: cgcgagctcagtcaaacag; reverse: aagttccttctggagcctctt), and *Stat5b* (forward: ggactccgtccttgataccg; reverse: tccatcgtgtcttccagatcg). Data were normalized to the *Actb* (β-actin) and *Ppia* expression. Relative quantification of mRNA was calculated by 2^-ΔΔCt^.

### Statistical analysis

When comparing two groups, the data were analyzed using a two-tailed, unpaired Student’s t-test. When comparing three groups simultaneously, one-way ANOVA followed by the Newman-Keuls multiple comparisons test was used. Two-way ANOVA and Holm–Sidak’s multiple comparisons test were used to analyze two variables (pregnancy and mutation effects). Statistical analyses were conducted with Prism software (GraphPad, San Diego, CA). All results are presented as mean ± standard error of the mean.

## Acknowledgements

We thank Ana M.P. Campos for her technical assistance. This study was funded by Fundação de Amparo à Pesquisa do Estado de São Paulo (FAPESP/Brazil; grant numbers: 2016/20897-3 to FW; 2020/01318-8 and 2024/21641-9 to JD; 2021/03316-5 to DOG; 2023/11833-5 to LMMS and 2024/22859-8 to GOS) and National Institutes of Health (NIA grant number: R01AG059779 to JJK and EOL).

## Authors’ Contributions

**Gabriel O. de Souza:** Formal analysis; Investigation. **Willian O. dos Santos:** Formal analysis; Investigation. **Frederick Wasinski:** Formal analysis; Investigation. **Ligia M. M. de Sousa:** Investigation. **Andressa G. Amaral:** Investigation. **Daniela O. Gusmao:** Investigation. **Edward O. List:** Resources. **John J. Kopchick:** Resources. **Gimena Fernandez:** Investigation. **Mario Perelló:** Investigation. **Carla R. P. Oliveira:** Investigation. **Manuel H. Aguiar-Oliveira:** Investigation. **Jose Donato Jr:** Conceptualization, Data Curation, Writing - Original Draft, Project administration, Funding acquisition.

## References

Aguiar-Oliveira, MH; Bartke, A 2019. Growth hormone deficiency: Health and longevity. Endocr Rev 40:575–601. 10.1210/er.2018-00216.

Aguiar-Oliveira, MH; Salvatori, R 2021. Disruption of the ghrh receptor and its impact on children and adults: The itabaianinha syndrome. Rev Endocr Metab Disord 22:81–89. 10.1007/s11154-020-09591-4.

Amorim, AR; Rossner, S; Neovius, M; Lourenco, PM; Linne, Y 2007. Does excess pregnancy weight gain constitute a major risk for increasing long-term bmi? Obesity (Silver Spring) 15:1278–1286. 10.1038/oby.2007.149.

Andreoli, MF; Donato, J; Cakir, I; Perello, M 2019. Leptin resensitisation: A reversion of leptin-resistant states. J Endocrinol 241:R81–R96. 10.1530/JOE-18-0606.

Augustine, RA; Ladyman, SR; Grattan, DR 2008. From feeding one to feeding many: Hormone-induced changes in bodyweight homeostasis during pregnancy. J Physiol 586:387–397. 10.1113/jphysiol.2007.146316.

Bartke, A; Kopchick, JJ 2015. The forgotten lactogenic activity of growth hormone: Important implications for rodent studies. Endocrinology 156:1620–1622. 10.1210/en.2015-1097.

Bridges, RS 1978. Retention of rapid onset of maternal behavior during pregnancy in primiparous rats. Behav Biol 24:113–117.

Bridges, RS 2016. Long-term alterations in neural and endocrine processes induced by motherhood in mammals. Horm Behav 77:193–203. 10.1016/j.yhbeh.2015.09.001.

Caron, P; Broussaud, S; Bertherat, J; Borson-Chazot, F; Brue, T; Cortet-Rudelli, C; Chanson, P 2010. Acromegaly and pregnancy: A retrospective multicenter study of 59 pregnancies in 46 women. J Clin Endocrinol Metab 95:4680–4687. 10.1210/jc.2009-2331.

Caufriez, A; Frankenne, F; Hennen, G; Copinschi, G 1993. Regulation of maternal igf-i by placental gh in normal and abnormal human pregnancies. Am J Physiol 265:E572–577. 10.1152/ajpendo.1993.265.4.E572.

Davey, HW; Xie, T; McLachlan, MJ; Wilkins, RJ; Waxman, DJ; Grattan, DR 2001. Stat5b is required for gh-induced liver igf-i gene expression. Endocrinology 142:3836–3841. 10.1210/endo.142.9.8400.

de Sousa, LMM; Vicente, VAN; Donato, J, Jr. 2025a. Negative feedback loops and hormonal factors that regulate gh secretion. Endocrinology 166:bqaf139. 10.1210/endocr/bqaf139.

de Sousa, ME; Gusmao, DO; Dos Santos, WO; Moriya, HT; de Lima, FF; List, EO; Kopchick, JJ; Donato, J, Jr. 2024. Fasting and prolonged food restriction differentially affect gh secretion independently of gh receptor signaling in agrp neurons. J Neuroendocrinol 36:e13254. 10.1111/jne.13254.

de Sousa, ME; Sousa, LMM; List, EO; Kopchick, JJ; Yakar, S; Kineman, RD; Donato, J, Jr. 2025b. Low liver-derived igf-1 drives the alterations in growth hormone secretion in food-restricted male mice. Endocrinology 166:bqaf148. 10.1210/endocr/bqaf148.

Dehkhoda, F; Lee, CMM; Medina, J; Brooks, AJ 2018. The growth hormone receptor: Mechanism of receptor activation, cell signaling, and physiological aspects. Front Endocrinol (Lausanne) 9:35. 10.3389/fendo.2018.00035.

Donbaloglu, Z; Singin, B; Acar, S; Bedel, A; Barsal Cetiner, E; Aydin Behram, B; Parlak, M; Tuhan, H 2023. Evaluation of the growth response of children with growth hormone deficiency according to the peak growth hormone levels in provocation tests. Arch Pediatr 30:573–579. 10.1016/j.arcped.2023.08.005.

Dunn, J; Dunn, C; Habbu, R; Bohay, D; Anderson, J 2012. Effect of pregnancy and obesity on arch of foot. Orthop Surg 4:101–104. 10.1111/j.1757-7861.2012.00179.x.

Emons, J; Chagin, AS; Savendahl, L; Karperien, M; Wit, JM 2011. Mechanisms of growth plate maturation and epiphyseal fusion. Horm Res Paediatr 75:383–391. 10.1159/000327788.

Frazao, R; Cravo, RM; Donato, J, Jr.; Ratra, DV; Clegg, DJ; Elmquist, JK; Zigman, JM; Williams, KW; Elias, CF 2013. Shift in kiss1 cell activity requires estrogen receptor alpha. J Neurosci 33:2807–2820. 10.1523/JNEUROSCI.1610-12.2013.

Furigo, IC; Kim, KW; Nagaishi, VS; Ramos-Lobo, AM; de Alencar, A; Pedroso, JA; Metzger, M; Donato, J, Jr. 2014. Prolactin-sensitive neurons express estrogen receptor-alpha and depend on sex hormones for normal responsiveness to prolactin. Brain Res 1566:47–59. 10.1016/j.brainres.2014.04.018.

Furigo, IC; Teixeira, PDS; de Souza, GO; Couto, GCL; Romero, GG; Perello, M; Frazao, R; Elias, LL; Metzger, M; List, EO; Kopchick, JJ; Donato, J, Jr. 2019. Growth hormone regulates neuroendocrine responses to weight loss via agrp neurons. Nat Commun 10:662. 10.1038/s41467-019-08607-1.

Gaete, X; Ferrer-Rosende, P; Pereira, A; Mericq, V 2025. Post-menarcheal growth patterns in a contemporary cohort of latino girls. Horm Res Paediatr 98:66–74. 10.1159/000536506.

Gardstedt-Berghog, J; Niklasson, A; Sjoberg, A; Aronson, AS; Pivodic, A; Nierop, AFM; Albertsson-Wikland, K; Holmgren, A 2024. Timing of menarche and pubertal growth patterns using the qeps growth model. Front Pediatr 12:1438042. 10.3389/fped.2024.1438042.

Garratt, M; Try, H; Smiley, KO; Grattan, DR; Brooks, RC 2020. Mating in the absence of fertilization promotes a growth-reproduction versus lifespan trade-off in female mice. Proc Natl Acad Sci U S A 117:15748–15754. 10.1073/pnas.2003159117.

Gatford, KL; Muhlhausler, BS; Huang, L; Sim, PS; Roberts, CT; Velhuis, JD; Chen, C 2017. Rising maternal circulating gh during murine pregnancy suggests placental regulation. Endocr Connect 6:260–266. 10.1530/EC-17-0032.

Georgescu, T; Swart, JM; Grattan, DR; Brown, RSE 2021. The prolactin family of hormones as regulators of maternal mood and behavior. Front Glob Womens Health 2:767467. 10.3389/fgwh.2021.767467.

Ghanem, I; Rizkallah, M 2020. The impact of residual growth on deformity progression. Ann Transl Med 8:23. 10.21037/atm.2019.11.67.

Greenhalgh, CJ; Alexander, WS 2004. Suppressors of cytokine signalling and regulation of growth hormone action. Growth Horm IGF Res 14:200–206. 10.1016/j.ghir.2003.12.011.

Gustafson, P; Ladyman, SR; Brown, RSE 2019. Suppression of leptin transport into the brain contributes to leptin resistance during pregnancy in the mouse. Endocrinology 160:880–890. 10.1210/en.2018-01065.

Kaur, H; Muhlhausler, BS; Sim, PS; Page, AJ; Li, H; Nunez-Salces, M; Clarke, GS; Huang, L; Wilson, RL; Veldhuis, JD; Chen, C; Roberts, CT; Gatford, KL 2020. Pregnancy, but not dietary octanoic acid supplementation, stimulates the ghrelin-pituitary growth hormone axis in mice. J Endocrinol 245:327–342. 10.1530/JOE-20-0072.

Ladyman, SR; Augustine, RA; Grattan, DR 2010. Hormone interactions regulating energy balance during pregnancy. J Neuroendocrinol 22:805–817. 10.1111/j.1365-2826.2010.02017.x.

Ladyman, SR; Carter, KM; Gillett, ML; Aung, ZK; Grattan, DR 2021. A reduction in voluntary physical activity in early pregnancy in mice is mediated by prolactin. eLife 10. 10.7554/eLife.62260.

Ladyman, SR; Fieldwick, DM; Grattan, DR 2012. Suppression of leptin-induced hypothalamic jak/stat signalling and feeding response during pregnancy in the mouse. Reproduction 144:83–90. 10.1530/REP-12-0112.

Ladyman, SR; Khant Aung, Z; Grattan, DR 2018. Impact of pregnancy and lactation on the long-term regulation of energy balance in female mice. Endocrinology 159:2324–2336. 10.1210/en.2018-00057.

Larsen, CM; Grattan, DR 2012. Prolactin, neurogenesis, and maternal behaviors. Brain Behav Immun 26:201–209. 10.1016/j.bbi.2011.07.233.

Liao, S; Vickers, MH; Stanley, JL; Baker, PN; Perry, JK 2018. Human placental growth hormone variant in pathological pregnancies. Endocrinology 159:2186–2198. 10.1210/en.2018-00037.

List, EO; Berryman, DE; Funk, K; Jara, A; Kelder, B; Wang, F; Stout, MB; Zhi, X; Sun, L; White, TA; LeBrasseur, NK; Pirtskhalava, T; Tchkonia, T; Jensen, EA; Zhang, W; Masternak, MM; Kirkland, JL; Miller, RA; Bartke, A; Kopchick, JJ 2014. Liver-specific gh receptor gene-disrupted (lighrko) mice have decreased endocrine igf-i, increased local igf-i, and altered body size, body composition, and adipokine profiles. Endocrinology 155:1793–1805. 10.1210/en.2013-2086.

Lopez-Tello, J; Yong, HEJ; Sandovici, I; Dowsett, GKC; Christoforou, ER; Salazar-Petres, E; Boyland, R; Napso, T; Yeo, GSH; Lam, BYH; Constancia, M; Sferruzzi-Perri, AN 2023. Fetal manipulation of maternal metabolism is a critical function of the imprinted igf2 gene. Cell Metab 35:1195–1208 e1196. 10.1016/j.cmet.2023.06.007.

Martinez-Garcia, M; Paternina-Die, M; Barba-Muller, E; Martin de Blas, D; Beumala, L; Cortizo, R; Pozzobon, C; Marcos-Vidal, L; Fernandez-Pena, A; Picado, M; Belmonte-Padilla, E; Masso-Rodriguez, A; Ballesteros, A; Desco, M; Vilarroya, O; Hoekzema, E; Carmona, S 2021. Do pregnancy-induced brain changes reverse? The brain of a mother six years after parturition. Brain Sci 11. 10.3390/brainsci11020168.

Miquet, JG; Sotelo, AI; Bartke, A; Turyn, D 2005. Desensitization of the jak2/stat5 gh signaling pathway associated with increased cis protein content in liver of pregnant mice. Am J Physiol Endocrinol Metab 289:E600–607. 10.1152/ajpendo.00085.2005.

Mounzih, K; Qiu, J; Ewart-Toland, A; Chehab, FF 1998. Leptin is not necessary for gestation and parturition but regulates maternal nutrition via a leptin resistance state. Endocrinology 139:5259–5262. 10.1210/en.139.12.5259.

Peris-Sampedro, F; Stoltenborg, I; Le May, MV; Zigman, JM; Adan, RAH; Dickson, SL 2021. Genetic deletion of the ghrelin receptor (ghsr) impairs growth and blunts endocrine response to fasting in ghsr-ires-cre mice. Mol Metab 51:101223. 10.1016/j.molmet.2021.101223.

Persechini, ML; Gennero, I; Grunenwald, S; Vezzosi, D; Bennet, A; Caron, P 2015. Decreased igf-1 concentration during the first trimester of pregnancy in women with normal somatotroph function. Pituitary 18:461–464. 10.1007/s11102-014-0596-3.

Pritschet, L; Taylor, CM; Cossio, D; Faskowitz, J; Santander, T; Handwerker, DA; Grotzinger, H; Layher, E; Chrastil, ER; Jacobs, EG 2024. Neuroanatomical changes observed over the course of a human pregnancy. Nat Neurosci 27:2253–2260. 10.1038/s41593-024-01741-0.

Punt, LD; Kooijman, S; Mutsters, NJM; Yue, K; van der Kaay, DCM; van Tellingen, V; Bakker-van Waarde, WM; Boot, AM; van den Akker, ELT; van Boekholt, AA; de Groote, K; Kruijsen, AR; van Nieuwaal-van Maren, NHG; Woltering, MC; Heijligers, M; van der Heyden, JC; Bannink, EMN; Rinne, T; Hannema, SE; de Waal, WJ; Delemarre, LC; Rensen, PCN; de Bruin, C; van Duyvenvoorde, HA; Visser, JA; Delhanty, PJD; Losekoot, M; Wit, JM; Joustra, SD 2025. Loss-of-function ghsr variants are associated with short stature and low igf-i. J Clin Endocrinol Metab 110:e1303–e1314. 10.1210/clinem/dgaf010.

Rooney, BL; Schauberger, CW; Mathiason, MA 2005. Impact of perinatal weight change on long-term obesity and obesity-related illnesses. Obstet Gynecol 106:1349–1356. 10.1097/01.AOG.0000185480.09068.4a.

Salvatori, R; Hayashida, CY; Aguiar-Oliveira, MH; Phillips, JA, 3rd; Souza, AH; Gondo, RG; Toledo, SP; Conceicao, MM; Prince, M; Maheshwari, HG; Baumann, G; Levine, MA 1999. Familial dwarfism due to a novel mutation of the growth hormone-releasing hormone receptor gene. J Clin Endocrinol Metab 84:917–923. 10.1210/jcem.84.3.5599.

Savage, MO; Bang, P 2012. The variability of responses to growth hormone therapy in children with short stature. Indian journal of endocrinology and metabolism 16:S178–184. 10.4103/2230-8210.104034.

Scholl, TO; Hediger, ML; Cronk, CE; Schall, JI 1993. Maternal growth during pregnancy and lactation. Horm Res 39 Suppl 3:59–67. 10.1159/000182785.

Scholl, TO; Hediger, ML; Schall, JI; Ances, IG; Smith, WK 1995. Gestational weight gain, pregnancy outcome, and postpartum weight retention. Obstet Gynecol 86:423–427.

Spear, BA 2002. Adolescent growth and development. J Am Diet Assoc 102:S23–29. 10.1016/s0002-8223(02)90418-9.

Teixeira, PDS; Couto, GC; Furigo, IC; List, EO; Kopchick, JJ; Donato, J, Jr. 2019a. Central growth hormone action regulates metabolism during pregnancy. Am J Physiol Endocrinol Metab 317:E925–E940. 10.1152/ajpendo.00229.2019.

Teixeira, PDS; Ramos-Lobo, AM; Furigo, IC; Donato, J 2019b. Brain stat5 modulates long-term metabolic and epigenetic changes induced by pregnancy and lactation in female mice. Endocrinology 160:2903–2917. 10.1210/en.2019-00639.

Travers, MT; Madon, RJ; Vallance, AJ; Barber, MC 1990. Circulating concentrations and hepatic expression of igf-1 during pregnancy and lactation in the mouse. Biochem Soc Trans 18:1268. 10.1042/bst0181268.

Trujillo, ML; Spuch, C; Carro, E; Senaris, R 2011. Hyperphagia and central mechanisms for leptin resistance during pregnancy. Endocrinology 152:1355–1365. 10.1210/en.2010-0975.

Udy, GB; Towers, RP; Snell, RG; Wilkins, RJ; Park, SH; Ram, PA; Waxman, DJ; Davey, HW 1997. Requirement of stat5b for sexual dimorphism of body growth rates and liver gene expression. Proc Natl Acad Sci U S A 94:7239–7244. 10.1073/pnas.94.14.7239.

Venken, K; Schuit, F; Van Lommel, L; Tsukamoto, K; Kopchick, JJ; Coschigano, K; Ohlsson, C; Moverare, S; Boonen, S; Bouillon, R; Vanderschueren, D 2005. Growth without growth hormone receptor: Estradiol is a major growth hormone-independent regulator of hepatic igf-i synthesis. J Bone Miner Res 20:2138–2149. 10.1359/JBMR.050811.

Vidal, A; Zhang, Q; Medigue, C; Fabre, S; Clement, F 2012. Dynpeak: An algorithm for pulse detection and frequency analysis in hormonal time series. PLoS One 7:e39001. 10.1371/journal.pone.0039001.

Wasinski, F; Pedroso, JAB; Dos Santos, WO; Furigo, IC; Garcia-Galiano, D; Elias, CF; List, EO; Kopchick, JJ; Szawka, RE; Donato, J, Jr. 2020. Tyrosine hydroxylase neurons regulate growth hormone secretion via short-loop negative feedback. J Neurosci 40:4309–4322. 10.1523/JNEUROSCI.2531-19.2020.

Wasinski, F; Tavares, MR; Gusmao, DO; List, EO; Kopchick, JJ; Alves, GA; Frazao, R; Donato, J, Jr. 2023. Central growth hormone action regulates neuroglial and proinflammatory markers in the hypothalamus of male mice. Neurosci Lett 806:137236. 10.1016/j.neulet.2023.137236.

Wasinski, F; Teixeira, PDS; List, EO; Kopchick, JJ; Donato, J, Jr. 2022. Growth hormone receptor contributes to the activation of stat5 in the hypothalamus of pregnant mice. Neurosci Lett 770:136402. 10.1016/j.neulet.2021.136402.

Waxman, DJ; O’Connor, C 2006. Growth hormone regulation of sex-dependent liver gene expression. Mol Endocrinol 20:2613–2629. 10.1210/me.2006-0007.

Wetz, HH; Hentschel, J; Drerup, B; Kiesel, L; Osada, N; Veltmann, U 2006. [changes in shape and size of the foot during pregnancy]. Orthopade 35:1124, 1126–1130. 10.1007/s00132-006-1011-1.

Wit, JM; Deeb, A; Bin-Abbas, B; Al Mutair, A; Koledova, E; Savage, MO 2019. Achieving optimal short- and long-term responses to paediatric growth hormone therapy. J Clin Res Pediatr Endocrinol 11:329–340. 10.4274/jcrpe.galenos.2019.2019.0088.

Woodside, B; Budin, R; Wellman, MK; Abizaid, A 2012. Many mouths to feed: The control of food intake during lactation. Front Neuroendocrinol 33:301–314. 10.1016/j.yfrne.2012.09.002.

Xu, XQ; Chen, Y; Wang, YR; Hu, FH; Li, J; Chang, GY; Li, X; Wang, R; Ding, Y; Wang, XM 2026. Final height prediction of girls at menarche: A combined model using left hand and wrist bone age, knee radiomic scores, and clinical characteristics. World J Pediatr 22:129–141. 10.1007/s12519-025-01002-5.

Younis, S; Shimonty, A; Perez-Menendez, S; You, X; Sridevan, SM; Ben-Tahar, S; Andrews, H; Shefelbine, S; Andersson, L; Charles, JF 2026. Persistent postnatal igf2 expression alters adult skeletal architecture in igf2 (g/a) mice. Bone Rep 30:101939. 10.1016/j.bonr.2026.101939.

Zampieri, TT; Ramos-Lobo, AM; Furigo, IC; Pedroso, JA; Buonfiglio, DC; Donato, J, Jr. 2015. Socs3 deficiency in leptin receptor-expressing cells mitigates the development of pregnancy-induced metabolic changes. Mol Metab 4:237–245. 10.1016/j.molmet.2014.12.005.

Zhou, Y; Xu, BC; Maheshwari, HG; He, L; Reed, M; Lozykowski, M; Okada, S; Cataldo, L; Coschigamo, K; Wagner, TE; Baumann, G; Kopchick, JJ 1997. A mammalian model for laron syndrome produced by targeted disruption of the mouse growth hormone receptor/binding protein gene (the laron mouse). Proc Natl Acad Sci U S A 94:13215–13220. 10.1073/pnas.94.24.13215.

Zigman, JM; Nakano, Y; Coppari, R; Balthasar, N; Marcus, JN; Lee, CE; Jones, JE; Deysher, AE; Waxman, AR; White, RD; Williams, TD; Lachey, JL; Seeley, RJ; Lowell, BB; Elmquist, JK 2005. Mice lacking ghrelin receptors resist the development of diet-induced obesity. J Clin Invest 115:3564–3572.

